# The *Ralstonia solanacearum* effector RipAV targets Plant U-box proteins and induces proteasomal-dependent degradation of BIK1

**DOI:** 10.1101/2025.01.06.631554

**Authors:** Jose S. Rufian, Xin Liu, Yaru Wang, Javier Rueda-Blanco, Gang Yu, Javier Ruiz-Albert, Alberto P. Macho

## Abstract

Plants have developed a complex immune system to detect and respond to invading pathogens. A critical aspect of this defence relies on regulatory mechanisms that control the activation of immune responses, ensuring these are efficient yet do not compromise overall plant performance. Many pathogens secrete proteins called effectors to manipulate plant cellular functions, suppressing plant immunity and promoting infection. Here, we show that the bacterial effector RipAV, secreted by the soil-borne bacterial pathogen *Ralstonia solanacearum*, hijacks a major immune signalling hub to suppress immune responses. RipAV targets members of the plant-specific ubiquitin ligase (PUB) family and calcium-dependent protein kinase 28 (CPK28), which has been shown to phosphorylate a set of PUBs to enhance their activity and regulate the stability of the key immune regulator BIK1. RipAV association enhances the CPK28-mediated phosphorylation of PUBs, inducing the proteasome-mediated degradation of BIK1 and the suppression of immunity. Importantly, we found that RipAV is required for maintaining a low accumulation of BIK1 during a *R. solanacearum* infection. These findings provide new insights into the sophisticated strategies employed by pathogens to subvert plant immunity.

## Introduction

Plants are constantly exposed to biotic stresses caused by pathogens. Among these, bacterial pathogens pose a significant threat to crops, leading to substantial yield losses worldwide. One notable example is *Ralstonia solanacearum*, the causal agent of bacterial wilt disease, which infects a wide range of plant species, including economically important crops (Hayward, 1991; Mansfield et al., 2012). *R. solanacearum* is a soil-borne pathogen that invades plants through the roots, colonizing the xylem vessels, where it proliferates, eventually leading to the blockage of the xylem, plant wilting, and death. The pathogenicity of *R. solanacearum* relies primarily on its virulence factors, the most critical being the Type III Secretion System (T3SS), which injects effector proteins, termed Type III Effectors (T3Es), directly inside plant cells, where they suppress the plant immune system and other plant cellular functions to promote bacterial proliferation and the development of disease (Landry et al., 2020; Macho, 2016).

Plants may recognize pathogens through plasma-membrane localized Pattern Recognition Receptors (PRR), which detect pathogen-associated molecular patterns (PAMPs), such as the elicitor peptides flg22 and elf18, present in bacterial flagellin and elongation factor Tu, respectively (Gómez-Gómez and Boller, 2000; Zipfel et al., 2006). Recognition of PAMPs triggers a defense mechanism known as pattern (or PAMP)-triggered immunity (PTI), characterized by a cascade of molecular events including a burst of reactive oxygen species (ROS), phosphorylation of MAP kinases, calcium influx, callose deposition, and stomata closure (Kadota et al., 2014; Li et al., 2014; Thor et al., 2020; Tian et al., 2019; Zhang et al., 2010). These events and others collectively limit pathogen proliferation and restrict its spread within the plant. A key protein involved in PTI activation is the receptor-like cytoplasmic kinase BIK1 (BOTRYTIS-INDUCED KINASE 1) (Lu et al., 2010; Zhang *et al*., 2010). The cellular levels of BIK1 are tightly regulated, and its activity is controlled through phosphorylation and ubiquitination (Bai et al., 2023; Lu *et al*., 2010; Ma et al., 2020; Zhang *et al*., 2010). Upon PAMP recognition, PRR complexes hyper-phosphorylate BIK1, activating its kinase activity (Lu *et al*., 2010; Zhang *et al*., 2010). Thus, the phosphorylation status of BIK1 defines two distinct pools within the cell: an active, phosphorylated form, and an inactive, non-phosphorylated form (Wang et al., 2018).

Ubiquitination plays a critical role in regulating BIK1 activity and homeostasis. Monoubiquitination of BIK1, mediated by the E3 ubiquitin ligase RING-H2 FINGER A3A (RHA3A), is required for dissociating phosphorylated BIK1 from PRR complexes, thereby enabling its downstream signalling (Ma *et al*., 2020). Additionally, two RING-type ubiquitin ligases, RING DOMAIN LIGASE 1 and 2 (RGLG1 and RGLG2), monoubiquitinate the inactive form of BIK1, thereby stabilizing its non-phosphorylated state (Bai *et al*., 2023). Members of the PLANT U-BOX (PUB) protein family, defined by a ∼70 amino acid U-box domain comprising a central alpha helix and two prominent surface-exposed loop regions, further regulate BIK1 levels in contrasting ways. The close homologs PUB22/23 and PUB25/26 target the inactive form of BIK1 for degradation (Wang *et al*., 2018). PUB25/26 activity is regulated by CALCIUM-DEPENDENT PROTEIN KINASE 28 (CPK28), which phosphorylates PUB25/26 at a conserved threonine residue (T95) upon flg22 perception, enhancing their activity and promoting BIK1 polyubiquitination and degradation following PTI activation (Wang *et al*., 2018). Furthermore, PUB4 exerts a dual role in BIK1 regulation: under basal conditions, it promotes degradation of the inactive form, while upon PTI activation, it protects the active, phosphorylated form of BIK1 from degradation (Yu et al., 2022).

The pivotal role of BIK1 in plant immunity is further underscored by the diverse strategies pathogens employ to compromise its function. The fungal pathogen *Fusarium oxysporum* utilizes its effector Avr2 to interfere with BIK1 monoubiquitination, thereby altering its subcellular localization (Blekemolen et al., 2022). In bacterial pathogens, *Pseudomonas syringae* delivers the effector AvrPphB, which cleaves BIK1 and other PBL proteins, effectively suppressing their immune functions (Zhang *et al*., 2010). Similarly, in *R. solanacearum*, the effector RipAW appears to directly interact with BIK1, leading to its ubiquitination and subsequent degradation (Sun et al., 2023). Another effector from *R. solanacearum*, RipAC, impairs PUB4 activity, inhibiting the protective role of PUB4 on the active form of BIK1, while simultaneously promoting PUB4-mediated degradation of the inactive pool (Yu *et al*., 2022).

In this study, we investigated the function of the *R. solanacearum* effector RipAV, originating from the strain GMI1000. This effector is widely distributed among *R. solanacearum* strains from phylotype I, IIA, IIB, and III (Peeters et al., 2013; Sabbagh et al., 2019). RipAV localizes to the cell periphery when transiently expressed in *Nicotiana benthamiana* (Denne et al., 2021) and contributes to GMI1000 virulence in eggplant (Macho et al., 2010). Given the limited characterization of RipAV, we explored its potential role in suppressing plant immunity. Our findings demonstrated that RipAV suppresses PTI in Arabidopsis. Additionally, RipAV interacts with various members of the PUB protein family but does not appear to affect their E3-ligase activity *in vitro*. Moreover, we found that RipAV associates to CPK28 and induces the CPK28-mediated phosphorylation of the conserved T95 in PUB22/23/24/25/26, ultimately leading to a proteasomal-mediated degradation of the non-activated form of BIK1. Our study uncovers an additional layer in the complex regulation of BIK1-mediated immune signalling and its manipulation by a bacterial pathogen to suppress immune responses.

## Results

### RipAV suppresses pattern-triggered immunity

To dissect the virulence contribution of RipAV and assess whether it suppresses plant immunity, we first generated Arabidopsis stable transgenic lines expressing *ripAV* fused to a 3xFLAG tag under a dexamethasone (DEX)-inducible promoter (**Figure S1A and B**). In the RipAV*-*3xFLAG expressing lines, ROS production triggered by the bacterial PAMPs flg22 and elf18 was significantly reduced compared to control plants (both Col-0 WT and a DEX-inducible GFP-expressing line) (**Figure 1A and B**). However, *ripAV* expression did not affect flg22-induced MAPK phosphorylation (**Figure 1C**), an independent early PTI response. As a later PTI response, we assessed the growth of a *P. syringae* DC3000-derivative strain carrying a mutation in *hrcC* (Δ*hrcC*) (Ronald et al., 1992). HrcC is an essential structural component of the T3SS, and therefore the 11*hrcC* strain is unable to deliver T3Es to suppress defence signalling, consequently activating PTI upon its inoculation in plant tissues (Ronald et al., 1992). RipAV-expressing plants were more susceptible to Δ*hrcC*, and the level of susceptibility correlated with RipAV-3xFLAG accumulation (**Figures 1D and S1B**). These findings indicate that RipAV specifically suppresses certain PTI responses, enhancing bacterial growth.

**Figure 1.**
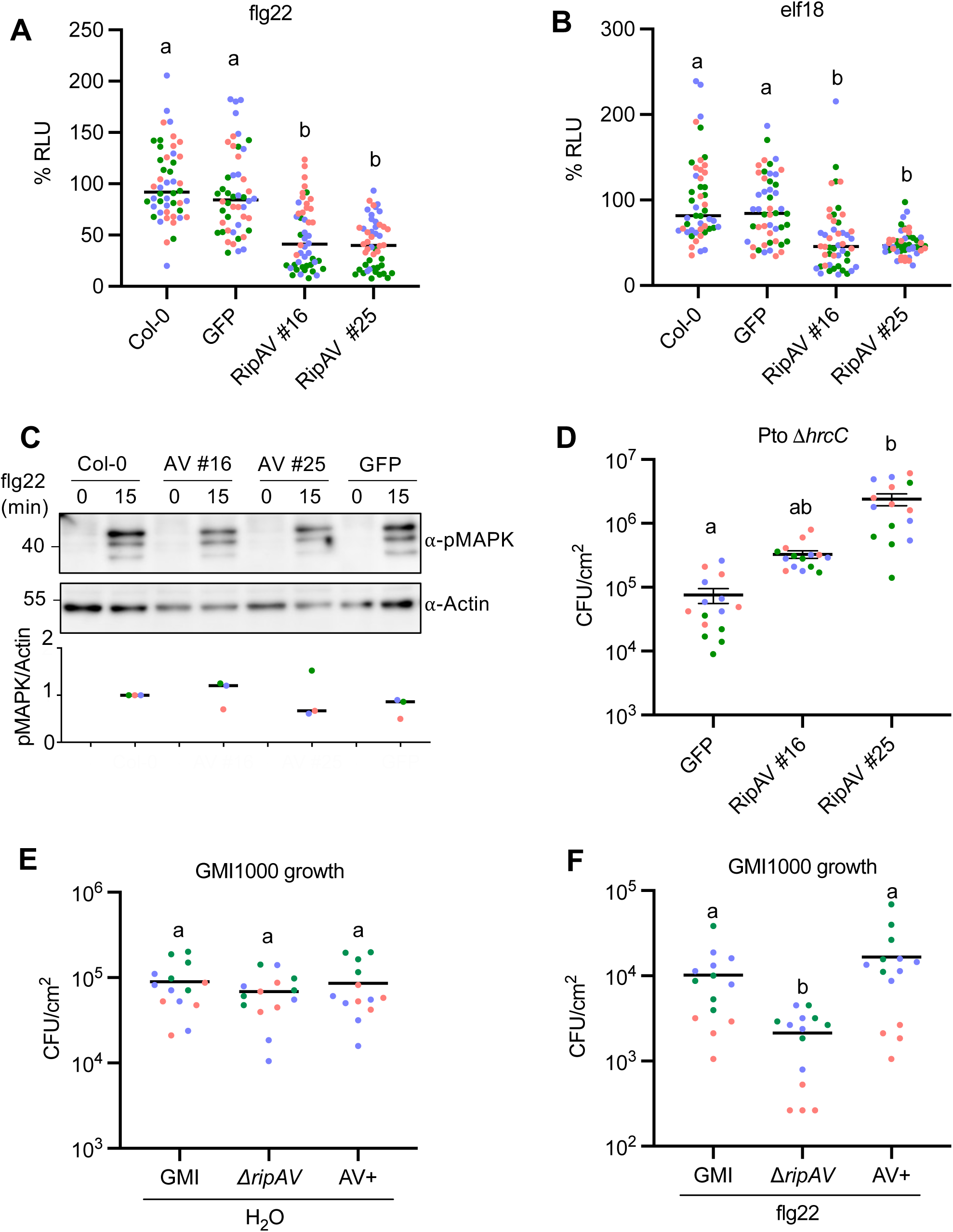
RipAV suppresses Pattern-Triggered Immunity. (**A,B**) RipAV suppresses ROS burst triggered by flg22 (**A**) and elf18 (**B**). Five-week old Arabidopsis leaf discs were immersed in 10 uM DEX for 24 h, followed by treatment with the different elicitors at 100 nM. ROS burst was analyzed using a luminol-based assay (n=16) and the experiment was repeated three times. Individual points represent percentage of Relative Luminiscense Units (RLU) in individual leaf discs, and different colors represent different replicates. (**C**) RipAV does not suppress flg22-induced phosphorylation of MAP Kinases. 10-days old Arabidopsis plants were immersed in 10 μM DEX for 24 h, followed by treatment with 1 μM flg22. Samples were taken both 0 and 15 minutes after treatment. The numbers below the blots indicate the ratios of the intensity of pMAPK into Actin. The experiment was repeated three times with similar results. (**D**) RipAV expression promotes the growth of *P. syringae* 11*hrcC*. Five-week old Arabidopsis leaves were infiltrated with 10 μM DEX. Three hours later, same leaves were syringe-infiltrated with a 10^5^ CFU/ml suspension of *P. syringae* 11*hrcC*. Bacterial titers were quantified three days later. Different points represent individual values group in colors based on different replicates. (**E,F**) *R. solanacearum* GMI1000 requires RipAV to partially overcome flg22-induced resistance. Arabidopsis leaves were treated with either water or 1 μM flg22. 24 hour later, same leaves were infiltrated with the indicated Ralstonia derivative (GMI = GMI1000; 11*ripAV = ripAV* deletion mutant; AV+ = RipAV complementation strain) Bacterial titers were quantified three days later. Different points represent individual values group in colors based on different replicates. In all graphs, statistically differences were determined by ANOVA (α=0.05) with Tukey’s multiple comparison test and different letters indicate statistical significance.

To further investigate the role of RipAV in PTI suppression during a bacterial infection, we generated a Δ*ripAV* knockout mutant and a complementation strain (AV+) in the *R. solanacearum* GMI1000 background (**Figure S1C**). We then performed a flg22-induced resistance assay followed by *Ralstonia* inoculation. While the growth of the 11*ripAV* mutant strain in Arabidopsis leaves was unaffected (**Figure 1E**), its growth was significantly impaired in plants pre-treated with flg22, but restored in the AV+ complementation strain (**Figure 1F**). Altogether, these results demonstrate that RipAV suppresses specific PTI responses and plays an important role in suppressing plant immunity during *Ralstonia* infection.

### RipAV interacts with Plant-U-box E3 ligases

To elucidate the function of RipAV in plant cells, we conducted a Yeast-Two-Hybrid screen using the N-terminal 400 amino acids of RipAV as bait and a library of cDNA from tomato roots inoculated with *R. solanacearum*. We identified three independent clones corresponding to the closest tomato ortholog of Arabidopsis PLANT U-BOX PROTEIN 22 (*AtPUB22*), which we named *SlPUB22* for consistency (**Figure 2A**). AtPUB22, along with its homologs AtPUB23 and AtPUB24, has been shown to participate in the regulation of plant immunity (Trujillo et al., 2008). Co-expression in *N. benthamiana* of RipAV-3xFLAG with AtPUB22, AtPUB23, and AtPUB24, each fused to a Green Fluorescent Protein (GFP) tag, followed by co-immunoprecipitation (Co-IP) confirmed that RipAV associates with AtPUB22, and revealed its association with AtPUB23 and AtPUB24 (**Figure 2B**). As the U-box domain was present in all the Yeast-Two-Hybrid clones (**Figure 2A**), we hypothesized that RipAV might associate with the U-box domain itself. Co-IP experiments with the U-box domain fused to GFP confirmed this interaction (**Figure 2B**). Further exploration showed that RipAV similarly interacts with additional PUB proteins, specifically AtPUB25 and AtPUB26, which are known to regulate plant immunity through BIK degradation (Wang *et al*., 2018) (**Figure 2C**). We additionally used Förster Resonance Energy Transfer-Fluorescence Lifetime Imaging (FRET-FLIM) to confirm the direct interaction between RipAV and all the tested PUB proteins (**Figure 2D**).

**Figure 2.**
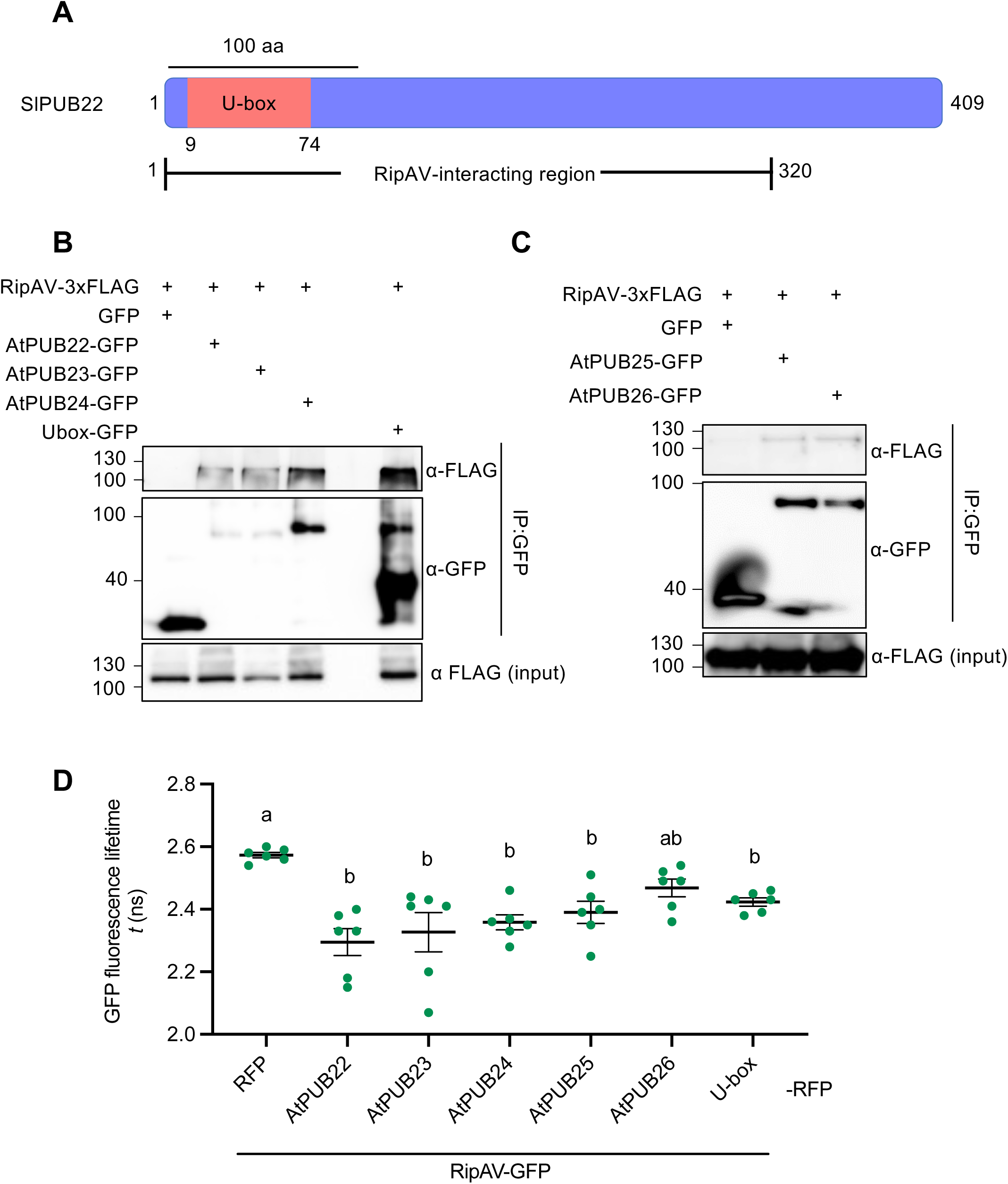
RipAV targets Plant U-box E3 ligases (PUB) proteins. (**A**) Schematic representation of the structure of tomato PUB22 and the RipAV interacting region found in different clones of a Yeast-Two-Hybrid screen. (**B,C**) Co-Immunoprecipitation of RipAV with Arabidopsis PUB22, PUB23, PUB24 (**B**) PUB25 and PUB26 (**C**). The *Agrobacterium* strains encoding the indicated PUBs (fused to GFP) were co-inoculated with RipAV-3xFLAG in *N. benthamiana*. Free GFP was used as negative control. 30 hours after inoculation, same leaves were treated with 10 uM DEX. Samples were taken 6 hours after DEX treatment. (**D**) Direct interaction between RipAV and PUBs was determined by FRET-FLIM. RipAV-GFP was co-expressed with the indicated proteins fused to RFP. Different point represents the quantification of the RipAV-GFP fluorescence lifetime. The experiment was repeated three times. Statistically differences were determined by ANOVA (α=0.05) with Tukey’s multiple comparison test and different letters indicate statistical significance.

### RipAV induces specific phosphorylation of PUBs

To further characterize the effect of RipAV on PUB proteins, we conducted *in vitro* ubiquitination assays using GST-fused SlPUB22, AtPUB22, AtPUB23, and AtPUB24 incubated either with MBP-RipAV or MBP-GFP (as negative control). RipAV did not alter the ubiquitin ladder resulting from PUB activity, suggesting no direct impact of RipAV on PUB E3-ligase activity *in vitro* (**Figure S2**).

Given the crucial role of phosphorylation in the activation of AtPUB25 and AtPUB26 (Wang *et al*., 2018), we examined whether their phosphorylation level is modified in the presence of RipAV. Arabidopsis protoplasts were co-transfected with FLAG-tagged AtPUB25 or AtPUB26 and either RipAV or an empty vector. Co-transfection with RipAV resulted in a general decrease in PUB protein accumulation (**Figure 3A**). However, this reduction was not specific to PUB proteins and also occurred when RipAV was co-expressed with GUS or GFP in Agrobacterium-mediated transient expression or in Arabidopsis protoplasts, respectively (**Figure S3A and B**). Interestingly, we observed an increased ratio of phosphorylated PUB proteins when co-expressed with RipAV (**Figure 3A**), suggesting that RipAV promotes PUB phosphorylation.

**Figure 3.**
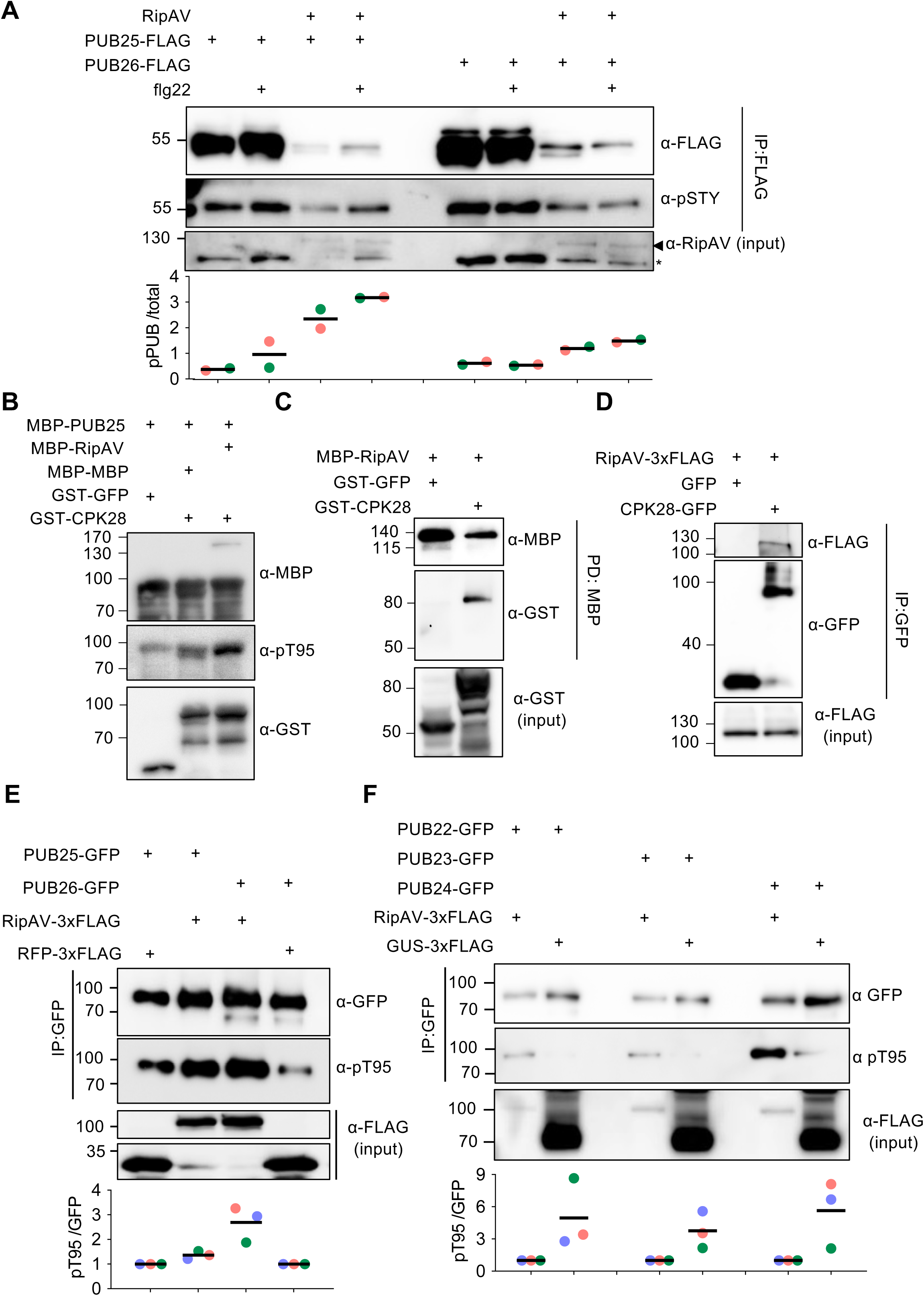
RipAV enhances phosphorylation of PUBs through association with CPK28. **(A)** Arabidopsis protoplasts were transfected with either PUB25 or PUB26 fused to FLAG and with a vector containing RipAV or an empty vector. Protoplasts were treated either with water or with 1 μM flg22 for 10 minutes, and a IP using FLAG beads was performed. Protein abundance was detected with western blot using the indicated antibodies and quantified with Fiji and represented as the ratio of phosphorylated PUB and total PUB. Asterisk indicate unspecific bands. The experiment was repeated twice with similar results. (B) Proteins were expressed in *E. coli* and purified accordingly with their respective tag. 200 ng of GST-CPK28 or GST-GFP was incubated with 2 ug of MBP-PUB25 and 1 μg of MPB-RipAV for 30 minutes at 30°C. Phosphorylation was detected with the specific pT95 antibody. The experiment was repeated twice with similar results. (**C**) MBP-RipAV was immobilized in an amylose resin and incubated with either GST-CPK28 or GST-GFP for 1 hour. Resin was washed and proteins were subjected to western blot using the indicated antibodies. (**D**) The indicated constructs were *Agrobacterium*-expressed in *N. benthamiana*. GFP-tagged proteins were immunoprecipitated, and association to RipAV-3xFLAG was detected with the appropriated antibodies. The experiment was repeated three times with similar results. (**E,F**) The indicated proteins were *Agrobacterium*-mediated transiently co-expressed in *N. benthamiana*. GFP-tagged proteins were immunoprecipitated and phosphorylation was detected with the specific antibody. Intensity of the bands was quantified with the Fiji software. The experiments were repeated three times with the results from the quantifications shown in the graphs at the bottom.

As phosphorylation by CPK28 of a conserved threonine residue (T95) in AtPUB25 increases its ubiquitin ligase activity (Wang *et al*., 2018), we hypothesized that RipAV may enhance this phosphorylation. *In vitro* kinase assays followed by immunoblot using an antibody that specifically recognizes phosphorylated T95 (Wang et al., 2018) showed that RipAV enhances CPK28-dependent T95 phosphorylation when co-incubated with AtPUB25 and CPK28 (**Figure 3B**), indicating that RipAV direcly enhances AtPUB25 phosphorylation by CPK28. This is supported by our observations that RipAV interacts directly with CPK28 as confirmed by in *in vitro* pull-down assays (**Figure 3C**) and that both proteins also associate *in planta* (**Figure 3D**). To assess *in planta* phosphorylation, we co-expressed RipAV-3xFLAG with AtPUB25 and AtPUB26 fused to GFP in *N. benthamiana*. After immunoprecipitation of GFP-tagged proteins, the ratio of T95-phosphorylated PUB proteins was significantly enhanced in the presence of RipAV (**Figure 3E**). Similar phosphorylation patterns were observed for AtPUB22, AtPUB23, and AtPUB24 (**Figure 3F**), demonstrating that RipAV enhances CPK28-mediated phosphorylation of multiple PUB proteins at T95.

### RipAV promotes proteasomal-dependent degradation of BIK1

Phosphorylation AtPUB25 and AtPUB26 at T95 enhances their E3 ubiquitin ligase activity, targeting BIK1 for degradation (Wang *et al*., 2018). Given that BIK1 is a central hub in plant immunity, we hypothesized that RipAV promotes BIK1 degradation. To test this hypothesis, we crossed the DEX-inducible *RipAV*-expressing line (#16) with an Arabidopsis transgenic line constitutively expressing BIK1-HA (Zhang *et al*., 2010), and quantified BIK1 accumulation. Upon DEX treatment, RipAV expression reduced BIK1 accumulation, which could be partially restored by treatment with the proteasome inhibitor MG132 (**Figure 4A**), indicating that RipAV promotes proteasomal-dependent degradation of BIK1. To study RipAV-induced degradation of BIK1 during a natural *Ralstonia* infection, we inoculated Arabidopsis plants expressing BIK1-HA with several *Ralstonia* strains, including the wild-type (GMI1000), a non-pathogenic strain carrying a knock-out mutation on the TTSS regulator gene *hrpG* (Δ*hrpG*) (Brito et al., 1999), the 11*ripAV* mutant, and its complemented strain. The results show that BIK1-HA accumulation in roots inoculated with the Δ*ripAV* mutant strain was higher in comparison to roots inoculated with the other strains analyzed (**Figure 4B and 4C**). Taken together, findings shown in Figure 4 indicate that RipAV induces proteasome-mediated BIK1 degradation during *Ralstonia* infection.

**Figure 4.**
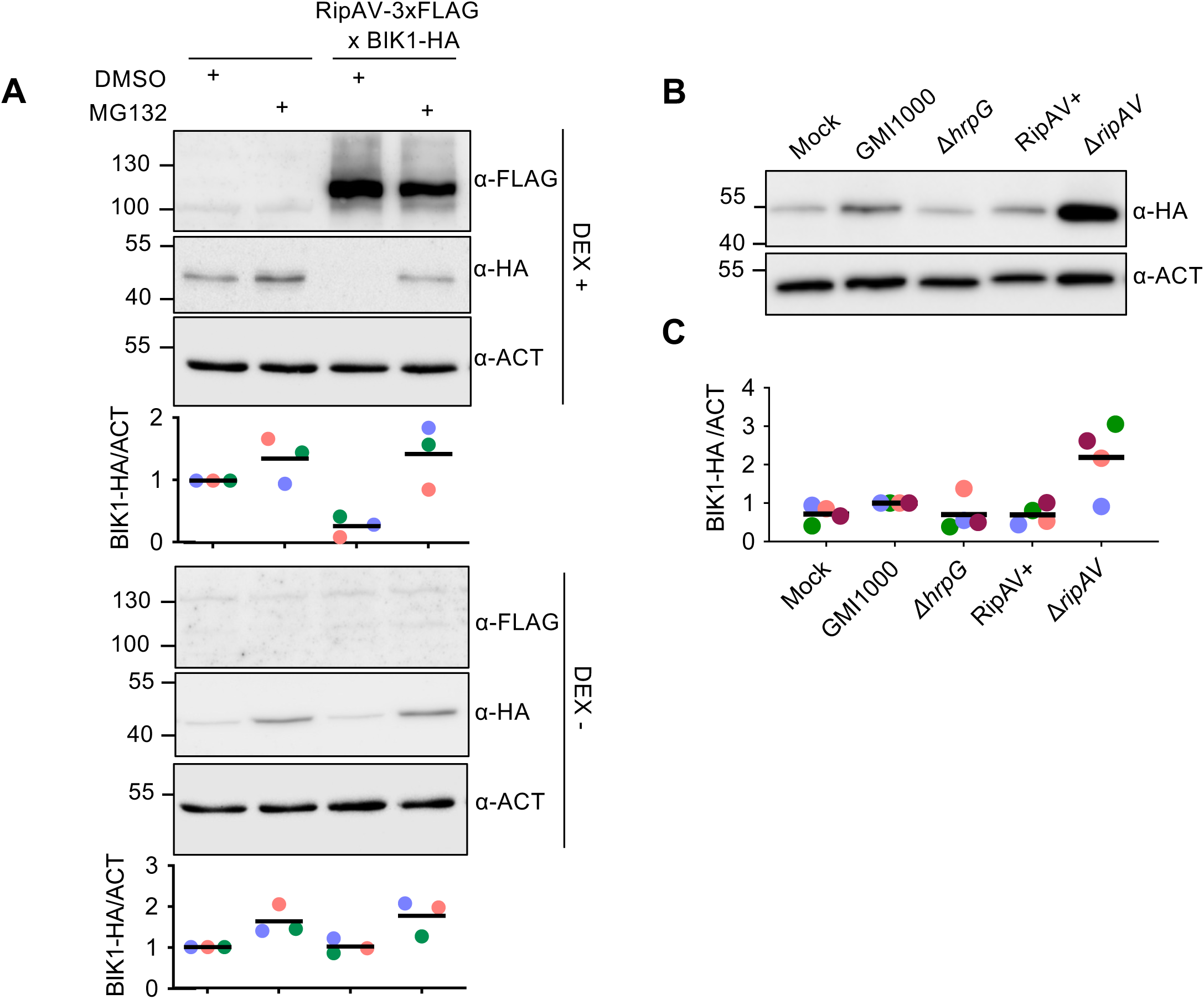
RipAV induces proteasome degradation of BIK1. (**A**) 4-week-old Arabidopsis leaves were treated with DEX to induce RipAV expression and either DMSO or 10 μM MG132. Samples were taken 6 hours after treatment. Levels of BIK1-HA were determined by Western blot. (**B**) Ten-days-old Arabidopsis seedlings were inoculated with 5 μl of a bacterial suspension corresponding to the strain indicated in the figure. 24 hours after infection, roots were detached and proteins extracted and analyzed by western blot. (**C**) Intensity of the bands was quantified with the Fiji software. The experiment was repeated three times with the results from the quantifications shown in the graphs at the bottom.

### RipAV contribution to *Ralstonia* virulence is partially redundant to that of RipAC

The reduction of non-activated BIK1 accumulation is expected to enhance bacterial proliferation by decreasing PTI signaling, and indeed we have shown the positive contribution of RipAV to GMI1000 replication under PTI-activating conditions (**Figure 1F**). To assess RipAV contribution to GMI1000 virulence in soil-drenching assays, we inoculated Arabidopsis plants with the Δ*ripAV* mutant strain and monitored disease progression. Consistent with our previous assays upon leaf infiltration (**Figure 1E**), the 11*ripAV* mutant did not exhibit a significant difference in virulence compared to the wild-type GMI1000 (**Figure 5A and 5B**). Bacterial T3Es are known to be functionally redundant, with a single *Ralstonia* strain secreting more than 70 T3Es (Peeters et al, 2013; Sabbagh et al, 2019). It is noteworthy that *Ralstonia* GMI1000 secretes another T3E, RipAC, which also affects the accumulation non-activated BIK1 through a different mechanism based on the manipulation of the E3-ligase PUB4 (Yu et al., 2022). Therefore, we hypothesized a functional redundancy between RipAC and RipAV through their manipulation of BIK1 accumulation. Indeed, while the Δ*ripAC* single mutant shows significantly attenuated virulence (**Figure 5**; Yu et al., 2020), a 11*ripAC/*11*ripAV* double mutant strain exhibited enhanced attenuation at later infection stages, resulting in increased plant survival (**Figures 5 and S4**). These results suggest partial functional redundancy between RipAV and RipAC.

**Figure 5.**
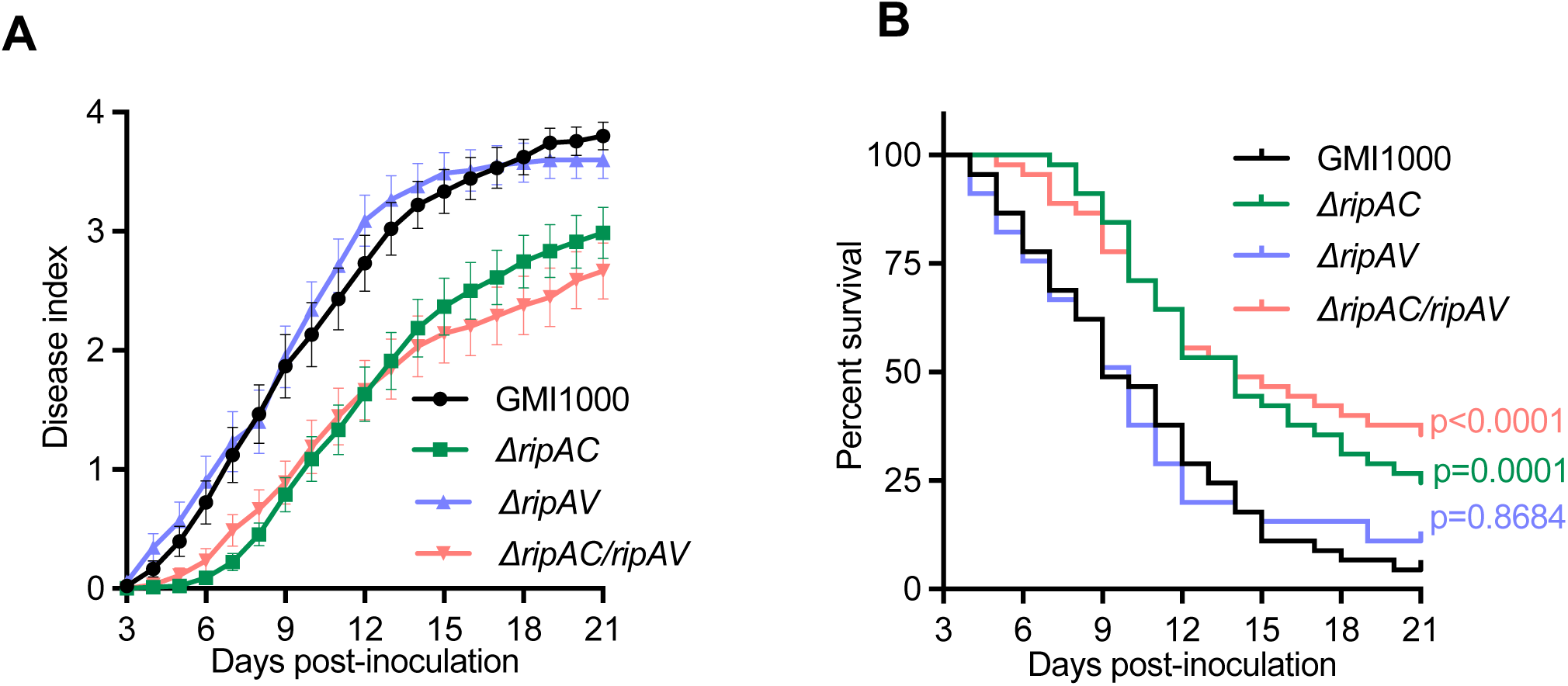
RipAV and RipAC are partially redundant. (**A**) Soil-drenching inoculation assays in Arabidopsis were performed with the indicated strains. n=15 plants per genotype. The results are represented as disease progression, showing the average wilting symptoms in a scale from 0 to 4 (mean ± SEM), (B) Survival curves of plants inoculated in (A). The disease score was transformed into binary data with the following criteria: a disease index lower than 2 was defined as “0” while a disease index equal to or higher than 2 was defined as “1” for each specific time point. Statistical analysis was performed using a log-rank (Mantel–Cox) test, and the corresponding *p* value is shown in the graph with the same color as each curve.

## Discussion

The Plant U-box (PUB) E3 ligases protein family has emerged as a key regulator of various cellular processes. Arabidopsis encodes a diverse repertoire of PUB proteins, comprising 64 members with variations in domain composition (Azevedo et al., 2001; Wiborg et al., 2008; Yee and Goring, 2009). PUB proteins have been implicated in processes such as hormone regulation, responses to abiotic stresses, and plant immunity regulation (Adler et al., 2017; Cho et al., 2008; Lu et al., 2011; Seo et al., 2016; Trujillo *et al*., 2008; Wang *et al*., 2018; Wang et al., 2022; Yi et al., 2024; Yu *et al*., 2022). Their significance in plant immune signalling is underscored by the number and diversity of pathogens that secrete effectors targeting PUB proteins to alter their functions. For instance, the effector AI106 from the insect *Apolygus lucorum* interacts with PUB33, a positive regulator of plant immunity; inhibition of PUB33 activity suppresses plant immunity and facilitates insect feeding (Dong et al., 2023). Similarly, the effector VDAL from the biotrophic fungal pathogen *Verticillium dahliae* interacts with PUB25/26, interfering with the association with their target, the transcription factor MYB6 (Ma et al., 2021). Reduced degradation of MYB6 enhances resistance to *V. dahliae* infection, potentially extending the biotrophic phase of infection (Ma *et al*., 2021). Additionally, the effector AVR3a from the oomycete *Phytophthora infestans* stabilizes the potato E3 U-box protein CMPG1, preventing host cell death during the biotrophic phase (Bos et al., 2010). Another example is the *Xanthomonas oryzae* effector XopP, which targets rice OsPUB44, a positive regulator of plant immunity; XopP specifically interacts with the U-box and linker domains of OsPUB44, inhibiting its activity and suppressing PTI (Ishikawa et al., 2014). Furthermore, the *R. solanacearum* effector RipAC targets PUB4, altering its phosphorylation status to direct the degradation of nonactive BIK1 (Yu *et al*., 2022). Collectively, these findings highlight the pivotal role of PUB proteins as key targets of pathogen effectors, emphasizing their critical function in modulating plant immunity.

In this study, we describe a mechanism by which the *R. solanacearum* effector RipAV suppresses PTI. RipAV hijacks a regulatory system of plant cells that fine-tune the immune response by degrading the non-activated pool of BIK1. Before PAMP perception, PUB25/26 maintains a basal activity that catalyses the polyubiquitination of BIK1, keeping its levels rate-limiting. Upon PAMP perception, CPK28 phosphorylation of PUB25^T95^/26^T94^ enhances their ubiquitin ligase activity, reducing nonactive BIK1 levels to regulate immune reactivation (Wang *et al*., 2018). Our results suggest that RipAV manipulates this system by promoting the phosphorylation of PUB22/23/24/25 at their respective PUB25^T95^-like residues (Figure 3), thereby reducing BIK1 levels and interfering with further immune signalling activation (**Figure 1A, 1B, 1D, and 1F**). Interestingly, RipAV does not affect MAPK phosphorylation after flg22 (Figure 1C) perception, which correlates with the previous observation that this pathway is independent of BIK1 (Ranf et al., 2014). Given that CPK28 promotes PUB25^T95^/26^T94^ phosphorylation after flg22 perception (Wang *et al*., 2018), RipAV association with CPK28 (**Figure 3C and 3D**) suggests that it could be promoting phosphorylation of PUBs, either in the absence of PAMP signalling, or after PTI has been suppressed by the action of other effectors (Landry *et al*., 2020). Our results show that a *R. solanacearum* single knockout mutant on *ripAV* is impaired in the suppression of BIK1 accumulation during a natural infection (**Figure 4B and 4C**), demonstrating the role of RipAV in maintaining low BIK1 levels long after the onset of infection. The absence of higher BIK1 accumulation in roots inoculated with the Δ*hrpG* mutant (a T3SS-deficient strain) is not surprising, as this mutant does not secrete effectors and does not cause an active infection, leading only to a basal activation of the immune system.

Among Arabidopsis PUB proteins, PUB22, 23, 24, 25, and 26 belong to the Class IV, characterized for the presence of a U-box domain and a C-terminal Armadillo (ARM) repeats, which mediate protein-protein interactions (Trenner et al., 2022). Bioinformatic analysis shows that PUB22-26 form a unique sub-class based on the presence of five undefined conserved motifs (Liao et al., 2021). One of these motifs (motif 1: DFDLTPNHTLRRLIQEWCVANRSYGVERIPTPKPPADK), includes the PUB25^T95^ residue, phosphorylated by CPK28 in PUB25 and PUB26 (**Figure 3B and 3E**; **Wang et al., 2018**), which is conserved and phosphorylated in PUB22, PUB23, and PUB24 (**Figure 3F**). Our results suggest that RipAV exploits this conserved mechanism to activate these PUBs and promote BIK1 degradation. PUB22/23/24 have been identified as negative regulators of plant immunity, with a triple *pub22/23/24* mutant exhibiting enhanced PTI responses and reduced susceptibility to *P. syringae* (Trujillo *et al*., 2008). PUB22 ubiquitinates Exo70B2, a subunit of the exocyst complex, inducing its proteasomal degradation, which is enhanced after flg22 treatment (Stegmann et al., 2012). Exo70B2 is involved in delivering the PRR FLS2 to the plasma membrane (Wang et al., 2020), and Arabidopsis *exo70B2* mutants display impaired PTI responses and increased susceptibility to *P. syringae* (Stegmann *et al*., 2012; Wang *et al*., 2020). Notably, MPK3 phosphorylates PUB22 *in vitro* at two threonine residues, one of which (T88) corresponds to PUB25^T95^ (Furlan et al., 2017). A phosphomimetic mutation in both residues results in an enhanced polyubiquitination of Exo70B2 (Furlan *et al*., 2017). Thus, RipAV activity could potentially also enhance Exo70B2 polyubiquitination and degradation, although this hypothesis would require further investigation.

Previous research has shown that the *R. solanacearum* effector RipAC alters PUB4 phosphorylation to promote degradation of nonactive BIK1 (Yu *et al*., 2022). Given the similarities in the mode of action of RipAC and RipAV, we considered the possibility of functional redundancy between these effector activities, which is supported by the results shown in Figure 5, revealing no significant impact of a single Δ*ripAV* mutation in *R. solanacearum* virulence in Arabidopsis. However, our results also show that in Arabidopsis leaves, when PTI is pre-activated, the Δ*ripAV* mutant exhibits defective growth (**Figure 1F**). Thus, the contribution to virulence of RipAV in Arabidopsis may be detected in specific conditions where RipAV activity is particularly relevant.

Functional redundancy within the effector repertoires can also mask the relevance of individual effectors (Ghosh and O’Connor, 2017). In *R. solanacearum* GMI1000, RipAC affects the same pathway as RipAV by inducing the degradation of nonactive BIK1 (**Figure 4**; (Yu *et al*., 2022)). A single Δ*ripAC* mutant shows a significant reduction in virulence (**Figure 5A**; (Yu et al., 2020)). RipAC is a multifunctional protein, not only interfering with BIK1 levels via PUB4 (Yu *et al*., 2022), but also suppressing immunity preventing MAPK-mediated phosphorylation of SGT1 (Yu *et al*., 2020) and altering root development to facilitate *Ralstonia* infection (Yu et al., 2024). Thus, the redundancy between RipAV and RipAC is only partial, which explains the modest virulence defect in the double 11*ripAC*11*ripAV* mutant compared to Δ*ripAC* (**Figure 5A**). Notably, the double mutant exhibited a higher survival rate (35%) compared to the other strains (4% for GMI1000, 11% for Δ*ripAV* and 24% for Δ*ripAC*) (**Figures 5B and S4**). As shown in **Figure S4**, most plants surviving GMI1000 infection did not display wilting symptoms, indicating that early suppression of PTI by these effectors is critical in determining the plant survival to *R. solanacearum* infection.

The complexity and sophistication of plant immune regulation comes at the price of vulnerability of its regulatory components to the manipulation by pathogens. This study uncovers an additional layer in the complex regulation of BIK1-mediated immune signaling, and reveals a novel mechanism by which *R. solanacearum* targets this signaling hub to suppress immune responses.

## Materials and Methods

### Plant materials and growth conditions

Arabidopsis plants Col-0 plants and transgenic lines used in this study were grown under short-day conditions (10 h light/14 h darkness, 22°C, 100-150 μE m^-2^ s^-1^) or long-day conditions (16h light/8 h darkness, 22°C, 100-150 μE m^-2^ s^-1^) according to the purpose of the experiment.

For *R. solanacearum* soil-drenching assays, Arabidopsis plants were cultivated in jiffy pots (Jiffy International, Kristiansand, Norway) in an environmentally controlled growth chamber under short-day conditions. After *R. solanacearum* inoculation, plants were incubated in a growth chamber under controlled conditions (75% humidity, 12 h light at 27°C, and 12 h darkness at 26°C). *N. benthamiana* plants were grown in a growth room (16h light/8 h darkness, 22°C, 100-150 μE m^-2^ s^-1^).

### Generation of transgenic plants

To generate Arabidopsis transgenic plants expressing RipAV under a dexamethasone (DEX)-inducible promoter, the ORF of RipAV was PCR-amplified and cloned into pENTR-D TOPO (Invitrogen) and transferred to pTA7001-3xFLAG (Li et al., 2013). The resulting plasmid (pTA-RipAV) was transformed into Agrobacterium GV3101. Arabidopsis Col-0 plants were transformed using the floral dipping method (Clough and Bent, 1998). Transgenic Arabidopsis plants were selected using hygromycin (50 μg/mL) and verified using immunoblots with an anti-FLAG antibody (Abmart #M20008). Two independent stable homozygous T3 lines were used for the experiments.

### Bacterial strains and cultivation conditions

*R. solanacearum* GMI1000 (Salanoubat et al., 2002) and derivative strains (Δ*ripAV, RipAV+,* 11*ripAC and* 11*ripAC*11*ripAV* strains) were grown at 28°C in rich BG medium (Plener et al., 2010). *P. syringae* pv. *tomato* derivative strain Δ*hrcC* (Ronald *et al*., 1992) and *A. tumefaciens* GV3101 (Weidi Bio #AC1001) carrying different constructs were grown at 28°C in Lysogeny Broth (LB) medium (Bertani, 1951). Antibiotics were used when appropriate, at the following concentrations: 25 μg/mL kanamycin, 10 μg/mL tetracyclin, 25 μg/mL rifampicin, 50 μg/mL spectinomycin.

To generate the *R*. *solanacearum* Δ*ripAV* mutant, the *ripAV* open reading frame (ORF) was replaced by a kanamycin resistance gene using homologous recombination (Zumaquero et al., 2010). To generate the knockout vector, 800 bp regions upstream (zone A) and downstream (zone B) the coding region of RipAV were amplified using the Q5 High-Fidelity DNA Polymerase (NEB) and the corresponding primers (Table S1). The resulting amplicons were fused by PCR and cloned into pEASYBLUNT cloning kit (TransGen Biotech #CB101-01). The kanamycin resistant cassette flanked by two FRT sequences was PCR-amplified and cloned in a *Eco*R I restriction site between zones A and B, generating the knockout vector. To generate the complementation vector, a region containing the RipAV promoter plus ORF was PCR-amplified and cloned into pENTR-D TOPO (Invitrogen) and transferred to the pRCT vector (Monteiro et al., 2012) by LR reaction. The plasmids were transformed into *R. solanacearum* by natural transformation (Perrier et al., 2018) and double recombinants were selected in the appropriated antibiotics. Insertions were confirmed by PCR and the *in vitro* secretion of RipAV was analyzed using the method described by (Lonjon et al., 2018). The double 11*ripAC*11*ripAV* strain was generated by transforming the RipAV knockout vector into Δ*ripAC* mutant (Yu *et al*., 2020).

### Bacterial virulence assays

For *R. solanacearum* soil-drenching assays, 15 four- to five-week-old Arabidopsis plants grown in Jiffy pots were inoculated with 300 mL of a 10^8^ CFU/mL suspension of the corresponding strain. Visual scoring of plant symptoms according to a scale ranging from ‘0’ (no symptoms) to ‘4’ (complete wilting) was performed as previously described (Vailleau et al., 2007)

For the quantification of *R. solanacearum* growth in Arabidopsis leaves, four- to five-week-old plant leaves were infiltrated with either water or 1 μM flg22. 24 hour later, same leaves were syringe-infiltrated with a 10^5^ CFU/mL suspension of the appropriate *Ralstonia* strain. Bacterial titers were quantified three days later by grinding three 10-mm-diameter leaf discs per plant and plating in BG medium plates.

For BIK1-HA quantification in inoculated Arabidopsis, plants were grown in ½ MS plates without sucrose (Yu et al., 2023). 10-days old plants were inoculated with a 5 μl drop of a 10^5^ CFU/mL bacterial suspension. 48 hours after inoculation, roots were collected, frozen in liquid nitrogen, and tissue was disrupted using a TyssueLyser II (Qiagen). Proteins were extracted in 50 μl of extraction buffer (100 mM Tris-HCl pH 8, 150 mM NaCl, 10% glycerol, 2 mM DTT, 1% (v/v) protease inhibitor cocktail, 0.5% (v/v) NP40, 10 mM sodium molybdate, 10 mM sodium fluoride, 2 mM sodium orthovanadate, 10 μM MG-132) and protein levels were detected through western blot using anti-HA (Roche #12CA5) and anti-actin (Agrisera #AS13 2640) antibodies.

To quantify the growth of the *P. syringae* pv. *tomato* DC3000 derivative Δ*hrcC*, four- to five-week-old plant leaves were infiltrated with 10 μM dexamethasone (Sigma, #D4902). 24 hour later, same leaves were syringe-infiltrated with a 10^5^ CFU/mL suspension of the Δ*hrcC* strain. Bacterial titers were quantified three days later by grinding three 10-mm-diameter leaf discs per plant and plating in LB medium plates.

### Measurement of ROS and MAPK activation

The ROS production triggered by PAMPs was measured using 4-week-old Arabidopsis plants, as previously described (Sang and Macho, 2017). Briefly, plant leaf discs were collected 2 days post-infiltration and placed in 96-well plates containing 10 μM dexamethasone. ROS was elicited with 100 nM elf18^Rsol^ or flg22^Pto^. Luminescence was measured over 60 min using a microplate luminescence reader (Varioskan flash, Thermo Scientific, USA).

For MAPK activation assays, 12-day-old seedlings (5 days on 1/2 MS solid agar plates, then 7 days in 1/2 MS liquid medium) were treated with 10 μM dexamethasone. 24 hours later, the seedlings were treated with a 100 nM flg22^Pto^ elicitor solution. Samples (2 seedlings per treatment) were collected 0 and 15 min after flg22 treatment. Proteins were extracted and subjected to immunoblots with anti-pMAPK antibody (Cell Signaling, 4370S). Immunoblots were also analyzed using anti-actin antibody to verify equal loading.

### Transient expression

Agrobacterium strain GV3101 with different constructs were suspended in infiltration buffer (10 mM MgCl_2_, 10 mM MES pH 5.6, and 150 μM acetosyringone) to a final OD_600_ of 0.5 and infiltrated into the abaxial side of *N. benthamiana* leaves using a 1 mL needless syringe. Leaf samples were collected 2 days after infiltration for the different experiments.

For transient expression in protoplasts, leaves from four-week-old Arabidopsis plants were detached and lower epidermis was removed following the tape-sandwich protocol (Wu et al., 2009). Protoplasts were then extracted and transfected following the protocol by Yoo et al (Yoo et al., 2007).

### Co-immunoprecipitation

Co-immunoprecipitation assays were performed as previously described (Sang et al., 2018). Briefly, 500 mg of ground *N. benthamiana* leaves were resuspended in 1 ml protein extraction buffer (100 mM Tris-HCl pH 8, 150 mM NaCl, 10% glycerol, 2 mM DTT, 1% (v/v) protease inhibitor cocktail, 0.5% (v/v) NP40, 10 mM sodium molybdate, 10 mM sodium fluoride, 2 mM sodium orthovanadate, 10 μM MG-132). Supernatants were filtered through micro bio-spin chromatography columns (BioRad). The filtered extracts were incubated with 15 µl GFP-trap agarose beads (ChromoTek, Germany) at 4 °C for 1 hour, followed by 4 washes with extraction buffer. The resulting protein samples were incubated at 70°C for 10 minutes in SDS loading buffer and loaded in SDS-PAGE acrylamide gels for western blot analysis using anti-GFP (Abicode, M0802-3a) and anti-FLAG (Abmart #M20008) antibodies. Both antibodies were diluted 1:5000.

### In vitro assays

For pull down assays, MBP-RipAV was expressed from pDEST566 and GST-GFP/ GST-CKP28 were expressed from pDEST565. Recombinant proteins were expressed in *E. coli* BL21 (DE3) Rosetta cells (Novagen, Darmstadt, Germany) after induction with 0.1 mM IPTG at 20 °C. Proteins were extracted in extraction buffer (20 mM Tris-HCl pH 7.5, 200 mM NaCl, 1 mM EDTA, 1mM DTT) and MBP-RipAV was immobilized into an amylose resin (New England Biolabs, USA). The extracts containing either GST-GFP or GST-CKP28 were incubated with the MBP-RipAV bound to the amylose resin for 1 hour at 4°C, followed with 5 washes with extraction buffer. Proteins were eluted in extraction buffer containing 10 mM maltose.

For *in vitro* ubiquitination assays, MBP- and GST-tagged proteins were purified as described above. The ubiquitination assay was performed as previously described (Furlan and Trujillo, 2017). Briefly, 0.2 ug of E1, 1.2 ug of E2, 2 ug of each PUB and 2 ug of either MPB-GFP or MBP-RipAV were mixed in incubation buffer and incubated 1 hour at 30°C. After that incubation, 2 μg of HA-Ub was added to the reaction, and it was incubated for an additional hour at 30°C.

To detect phosphorylation of PUB25^T95^, MBP- and GST-tagged proteins were purified as described above, and the assay was performed as previously described (Wang *et al*., 2018). Briefly, 200 ng of either GST-GFP or GST-CPK28, 2 μg of MBP-PUB25 and 1 μg of either MBP-MBP or MBP-RipAV were incubated for 30 minutes at 30°C.

### FRET-FLIM

Förster resonance energy transfer-fluorescence lifetime imaging (FRET-FLIM) was performed as previously described (Yu *et al*., 2022). Briefly, RipAV-GFP was expressed from pGWB505 and the indicated PUBs fused to RFP were expressed from pGWB554. The experiment was performed using a Leica TCS SMD FLCS confocal microscope. Two days after infiltration*, N. benthamiana* leaves co-expressing the indicated proteins were observed under the microscope. Accumulation of the GFP- and RFP-tagged proteins was estimated before measuring lifetime. Samples were excited with the tuneable WLL set at 488 nm with a pulsed frequency of 40 MHz, and emission was detected using SMD GFP/RFP Filter Cube. The fluorescence lifetime shown in the figures corresponds to the average fluorescence lifetime of the donor collected and analyzed by PicoQuant SymphoTime software. Mean lifetimes are presented as mean +/- SEM based on eight images from three independent experiments.

### Generation of custom antibody

The custom polyclonal anti-RipAV antibody was generated in rabbits using a peptide containing the amino acids 1-400 of RipAV as antigen (Abclonal Co., China). The specificity of the anti-RipAV antibody was determined by immunoblot analyses using RipAV-3xFLAG transient expression in *N. benthamiana*.

### Statistical analysis

Statistical analyses were performed using GraphPad Prism 9. Unless stated otherwise, data are presented as mean ± SEM, and the methods for statistical analysis are described in the figure legends.

## Acknowledgements

We thank Rosa Lozano-Duran for critical reading of this manuscript, Xinyu Jian and Fangyuan Wu for technical and administrative assistance during this work, Jian-Min Zhou for sharing biological materials, and all the members of the Macho and Lozano-Duran laboratories for helpful discussions. We thank the PSC Cell Biology core facility for assistance with confocal microscopy. This work was supported by the Strategic Priority Research Program of the Chinese Academy of Sciences (grant XDB27040204 to A.P.M.), the Chinese 1000 Talents Program (to A.P.M.), the Shanghai Center for Plant Stress Biology (Center for Excellence in Molecular Plant Sciences, Chinese Academy of Sciences to A.P.M.), the Science and Technology Commission of Shanghai Municipality (grant 20WZ2503700 to J.S.R) and by I Plan Propio de Investigación y Transferencia (Universidad de Málaga, grant B1-2021_28 to J.S.R.). J.R.B was supported by a travel grant from University of Malaga (D.2) and Project Grant PID2021-127245OB-I00 financed by MCIN/AEI/10.13039/501100011033/ and ‘ERDP A way of making Europe’. The authors have no conflict of interest to declare.

## Author contributions

JSR and APM planned and designed the research. JSR performed most of the experiments. XL, YW, JR-B, and GY performed additional experiments. JR-A provided resources and advice for the project. JSR wrote the manuscript with inputs from all the authors.

## Data availability

All the data supporting the findings of this study are included in this article or as supporting information.

---

**Figure S1. Characterization of RipAV transgenic plants and Ralstonia mutant. (A)** 4-weeks-old Arabidopsis plants of the indicated genotypes grown in soil. (**B**) Three leaves of plants in (A) were infiltrated with 10 μM DEX or water. 24 hours later, tissue was collected and proteins extracted. RipAV-3xFLAG was detected by western blot. (**C**) RipAV detection in supernatants of the indicated genotypes. The effector was detected with a custom antibody (αRipAV)

**Figure S2. RipAV does not affect E3 activity of PUBs *in vitro*.** Purified proteins were mixed as indicated in the figure and incubated for 1 hour at 30°C. The in vitro ubiquitination assay was started after that incubation by the addition of 2 ug of HA-UBIQUITIN. The reaction was incubated for 1 hour at 30°C. Proteins were subjected to western blot and probed with the indicated antibody. Panels correspond to assays with SlPUB22 (A), AtPUB22 (B), AtPUB23 (C) and AtPUB24 (D).

**Figure S3. Co-expression with RipAV reduces protein accumulation. (A)** Transient expression in *N. benthamiana* of either RipAV-3xFLAG or GFP-3xFLAG and GUS-HA. (**B)** Protein accumulation after transfection in Arabidopsis protoplasts of either RipAV-HA or GUS-HA and GFP. Proteins were detected with the indicated antibodies.

**Figure S4. Photographs of plants surviving the soil-drenching experiments.** Plants were inoculated with the indicated strains and incubated for 21 days with constant watering.

