## Supplemental information for "The *Ralstonia solanacearum* effector RipAV targets Plant U-box proteins and induces proteasomal-dependent degradation of BIK1"

Figure S1

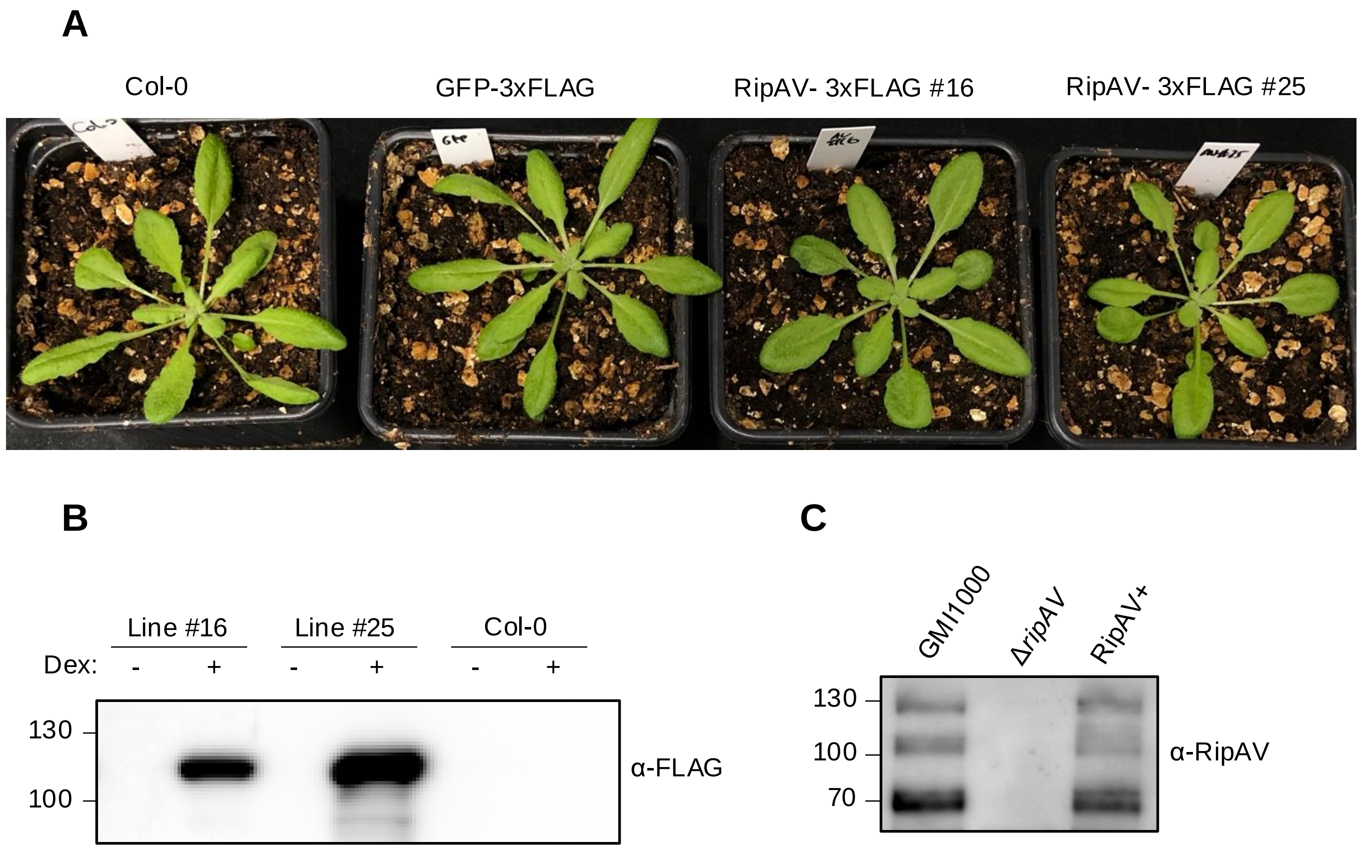


**Figure S1. Characterization of RipAV transgenic plants and *Ralstonia* mutant. (A)** 4-weeks-old Arabidopsis plants of the indicated genotypes grown in soil. (**B**) Three leaves of plants in (A) were infiltrated with 10 μM DEX or water. 24 hours later, tissue was collected and proteins extracted. RipAV-3xFLAG was detected by western blot. (**C**) RipAV detection in supernatants of the indicated genotypes. The effector was detected with a custom antibody (αRipAV)

### Figure S2

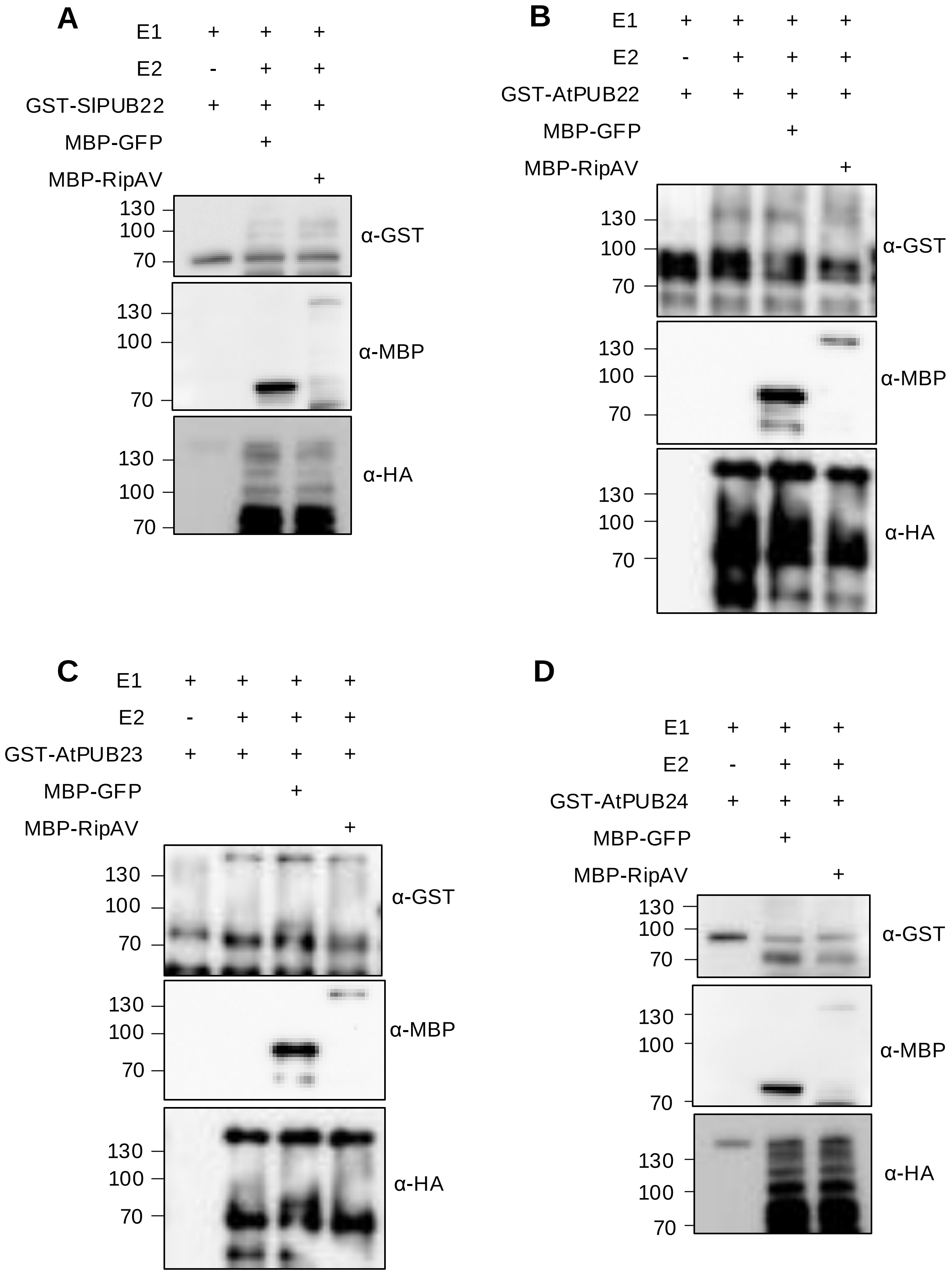


**Figure S2. RipAV does not affect E3 activity of PUBs *in vitro*.** Purified proteins were mixed as indicated in the figure and incubated for 1 hour at 30ºC. The in vitro ubiquitination assay was started after that incubation by the addition of 2 ug of HA-UBIQUITIN. The reaction was incubated for 1 hour at 30ºC. Proteins were subjected to western blot and probed with the indicated antibody. Panels correspond to assays with SlPUB22 (A), AtPUB22 (B), AtPUB23 (C) and AtPUB24 (D).

Figure S3

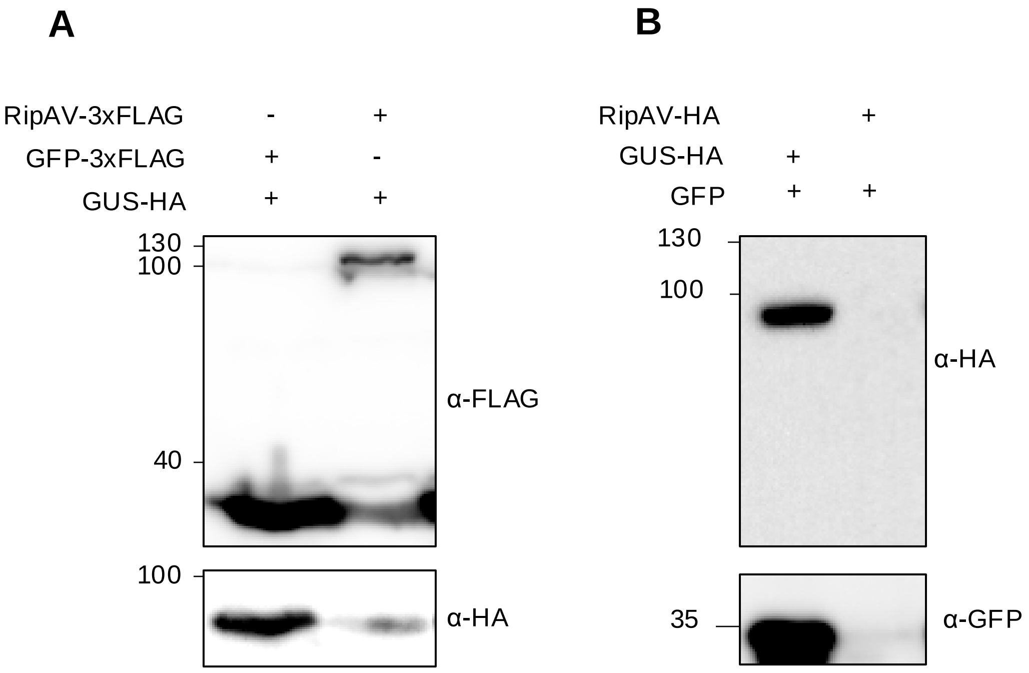


**Figure S3. Co-expression with RipAV reduces protein accumulation. (A)** Transient expression in *N. benthamiana* of either RipAV-3xFLAG or GFP-3xFLAG and GUS-HA. (**B)** Protein accumulation after transfection in Arabidopsis protoplasts of either RipAV-HA or GUS-HA and GFP. Proteins were detected with the indicated antibodies.

### Figure S4

GMI1000 | Δ*ripAC* | Δ*ripAV* | Δ*ripAC*/ Δ*ripAV*

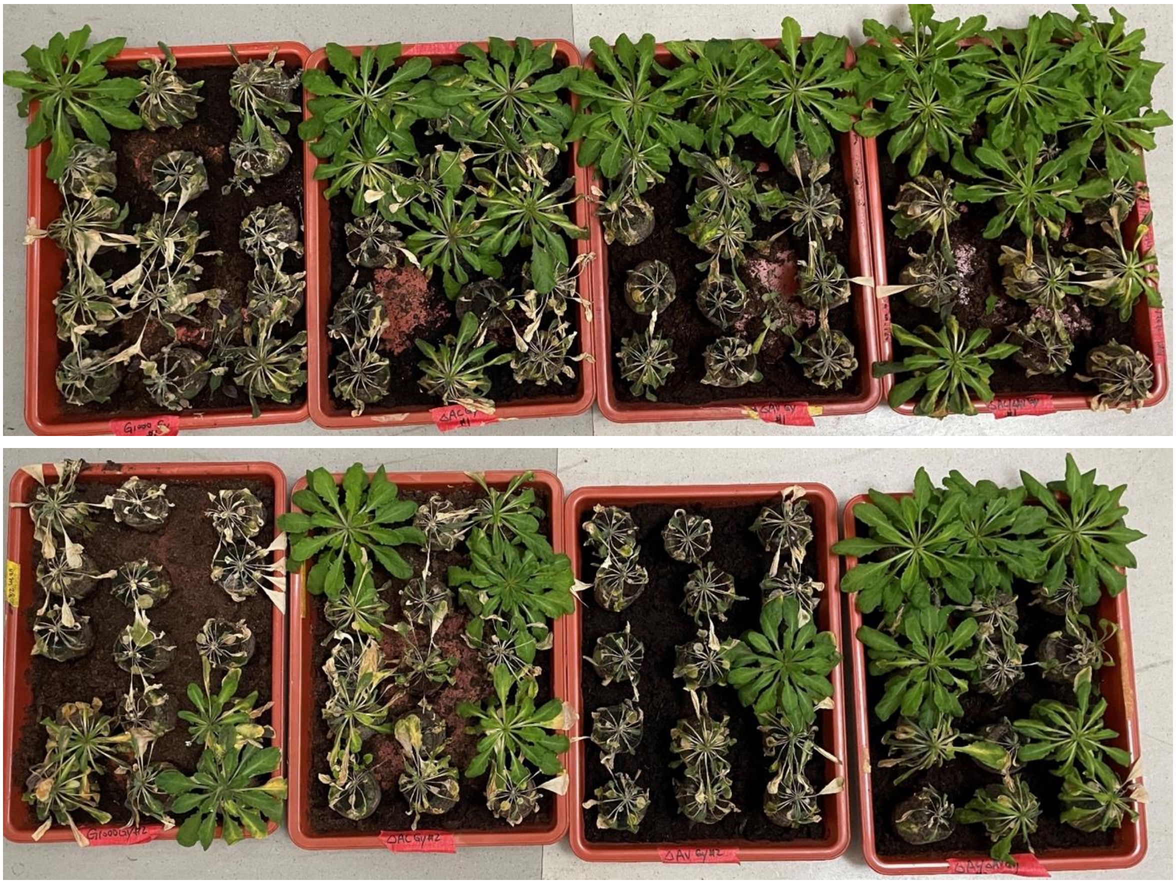

**Figure S4. Photographs of plants surviving the soil-drenching experiments.** Plants were inoculated with the indicated strains and incubated for 21 days with constant watering.
